# CRISPR/Cas9-mediated integration of large transgene into pig potential safe harbor

**DOI:** 10.1101/722348

**Authors:** Guoling Li, Xianwei Zhang, Haoqiang Wang, Jianxin Mo, Cuili Zhong, Junsong Shi, Rong Zhou, Zicong Li, Huaqiang Yang, Zhenfang Wu, Dewu liu

**Author notes:** Corresponding author: Zhenfang Wu and Dewu liu.

## Abstract

Clustered regularly interspaced short palindromic repeats/CRISPR-associated protein 9 (Cas9) is a precise genome manipulating tool which can produce targeted gene mutations in various cells and organisms. Although CRISPR/Cas9 can efficiently generate gene knock-out, the gene knock-in efficiency mediated by homology-directed repair (HDR) remains low, especially for large fragment integration. In this study, we established an efficient method for CRISPR/Cas9-mediated integration of large transgene cassette carrying salivary gland-expressing multiple digestion enzymes (≈ 20 kbp) in *CEP112* locus in pig fetal fibroblasts. Our results showed that using homologous donor with a short left arm and a long right arm yielded the best CRISPR/Cas9-mediated knock-in efficiency, and the targeting efficiency in potential safe harbor *CEP112* locus are higher than *ROSA26* locus. The *CEP112* knock-in cell lines were used as nuclear donors for somatic cell nuclear transfer to create genetically modified pigs. We found that knock-in pig (705) successfully expressed three microbial enzymes (β-glucanase, xylanase, and phytase) in salivary gland, suggesting potential safe harbor *CEP112* locus supports exogenous genes expression by a tissue-specific promoter. In summary, we successfully targeted *CEP112* locus using our optimal homology arm system for precise modification of pigs, and established a modified pig model for foreign digestion enzyme expression in saliva.

## 1. Introduction

Genetically modified animals play an important role in agricultural and biomedical studies. Zinc-finger nucleases (ZFNs), transcription activator like effector nucleases (TALENs) and bacterial clustered regularly interspaced short palindromic repeats (CRISPR)-associated protein 9 (Cas9) system can enable the targeted generation of DNA double strand breaks (DSBs) with a high accuracy and efficiency (Doudna and Charpentier 2014; Hsu *et al*. 2014). The induced DSBs trigger two major DNA repair systems, namely non-homologous end joining (NHEJ) and homology-directed repair (HDR) (You *et al*. 2009). The NHEJ pathway results in gene knockout (KO) by generating randomly sized small deletions or insertions (Indels) in the target gene, and HDR enables knock-in (KI) which represents a precise type of genome editing in the presence of homogenous template (Doudna and Charpentier 2014; Hsu *et al*. 2014). However, CRISPR/Cas9-mediated gene editing is quiet inefficient (Li *et al*. 2014; Richardson *et al*. 2016), especially in primary cells (Li *et al*. 2017a; Li *et al*. 2018). Meanwhile, the length of some transgenic cassettes are very long, such as mammary gland bioreactors vectors (>25 kbp), salivary gland bioreactor carriers (≈20 kbp) and multi-gene co-expression vectors. Therefore, precise insertion of large transgenic fragment in primary cells remains a big challenge.

Although numerous methods indicated that inhibiting the NHEJ pathway or activating the HDR pathway can efficiency promote KI efficiency, they mainly focus on immortal cell lines, and have the problem of cytotoxicity and integration fragments shorter than <2 kbp (Chu *et al*. 2015; Maruyama *et al*. 2016; Song *et al*. 2016). Recently, researchers reported that micro-homology-mediated end joining (MMEJ) can highly efficiently integrated large fragments (5.7 kbp to 9.6 kbp) by using 10 to 50 bp micro-homology (Sakuma *et al*. 2015; Sakuma *et al*. 2016; Nakamae *et al*. 2017), but there is still no success for targeted integration of large fragments over 20 kbp. Meanwhile, MMEJ system is more likely to lead to random integration. In addition, Yoshimi et al reported two-hit two-oligo with plasmid system (2H2OP) that simultaneously injected 2 CRISPR/Cas9 system, 2 strands 120 bp single-stranded oligonucleotide, and donor vector into the fertilized eggs, could integrate the large fragments of approximately 200 kbp into the genomic region with an efficiency of 1/15 (6.7%) (Yoshimi *et al*. 2016). However, the disadvantage of 2H2OP is the high rate of indel mutations at ssODN-mediated conjunction sites, which could impair the integrity of inserted transgene.

Pigs are important agricultural animals with economic values and also used as ideal human disease models (Yang *et al*. 2016; Zheng *et al*. 2017), but precise modifications for generation of KI pigs are always a tough work (Li *et al*. 2017; Li *et al*. 2018). In the previous study, we have successfully established transgenic (TG) pigs co-expressing three microbial enzymes (β-glucanase, xylanase, and phytase) in salivary gland (Zhang *et al*. 2018). These enzymes can degrade plant non-starch polysaccharides (NSP) and phytate. These TG pigs can significantly promote the digestion of nitrogen (N) and phosphorus (P) in formula feed, offering a promising approach to improve feed efficiency and reduce the impacts on the environment. By using piggybac-mediated transgenesis methods, we generated TG pig lines harboring the target gene in the intron of *CEP112* which encodes a centrosomal protein of 112 kDa and is localized around spindle poles, but its structure and function are still unclear (Jakobsen *et al*. 2011; Shashikala *et al*. 2013). The pig lines in our study can efficiently express the foreign gene that can be stably inherited to the offspring, and showed normal and healthy performance regardless of heterozygote or homozygote, which indicates *CEP112* has the potential as a safe site for the expression of foreign genes like the *ROSA26* locus that is frequently used for creating modified animals to achieve ubiquitous or conditional gene expression in vivo (Li *et al*. 2014; Xie *et al*. 2017). To further establish modified pigs from different pedigrees for breeding of new pig variety harboring the above-mentioned advantageous traits, we tried to insert the transgene cassette into the specific genomic locus, including *CEP112 or ROSA26*. Thus, in this study, we optimized CRISPR/Cas9-mediated KI strategy at *ROSA26* and *CEP112* locus in porcine foetal fibroblasts (PFFs) by using different sizes of homologous arms, and then generated modified pigs expressing β-glucanase, xylanase, and phytase by highly efficiently integrating transgene fragment (≈ 20 kbp) into potential safe harbor *CEP112* locus.

## 2. Materials and methods

### 2.1 Ethics statement

The animal use protocol was in accordance with the Instructive Notions with Respect to Caring for Laboratory Animals issued by the Ministry of Science and Technology of China. The use of animal was approved by the Institutional Animal Care and Use Committees in South China Agricultural University.

### 2.2 Plasmid construction

Cas9-gRNA plasmid PX330 was obtained from Addgene (Plasmid #42230). Seven *CEP112* sgRNAs were designed in http://crispr.mit.edu (**Supplementary Table S1**), and a *ROSA26* sgRNA (R5: 5’-GTGAGAGTTATCTGACCGTA-3’) was used as previous report (Li *et al*. 2017). The *CEP112* sgRNAs were synthesized, annealed, and cloned into PX330, to form the targeting plasmids PX330-C1 to PX330-C7. For the homologous template DNA of the *ROSA26* and *CEP112* locus, we amplified left arm (LA) and right arm (RA) on both sides of targeted sites (**Supplementary Table S2**), and constructed into the upstream and downstream regions of transgene pPB-mPSP-BgEgXyAp-neoGFP (PB-PSP-BEXA) to form a donor vector (Zhang *et al*. 2018). For the transgene, two β-glucanases genes (*bg17A* and *eg1314*), a xylanase gene (*xynB*) and a phytase gene (*appA*) were fused in a head-to-tail tandem array and E2A, P2A and T2A were linkers between them, abbreviated as BgEgXyAp gene.

### 2.3. Cell culture and transfection

PFFs were isolated from a 30-day old fetus of Duroc pig and cultured in DMEM (Thermo Fisher Scientific, Suwanee, GA,USA) supplemented with 10% fetal bovine serum (Thermo Fisher Scientific, Suwanee, GA,USA). To evaluate efficiency of induced-DSBs, PFFs were grown up to 80% confluence, harvested by 0.25% trypsin/EDTA and resuspended in 100 μL nucleofector solution (Amaxa Biosystems/Lonza, Cologne, Germany) containing 5 μg CRISPR/Cas9 plasmid, then electroporated by the program A-033 with Nucleofector 2b Device (Amaxa Biosystems/Lonza, Cologne, Germany). Afterward, the target region was amplified with primer shown in **Supplementary Table S3**, and PCR products were evaluated by T7 endonuclease I assay as previously described (Li *et al*. 2017). For KI cell lines screening, PEFs were co-electroporated with 5 μg CRISPR/Cas9 plasmid and 10 μg donor template. After electroporation, cells were divided into appropriate well plates for subsequent culture and screening.

### 2.4. Evaluation of KI cell lines by PCR amplification

One day after co-electroporation with CRISPR/Cas9 plasmid and donor template, cells were transferred into 10 cm plates, and selected with G418 (400 μg/ml) for approximately two weeks. The fluorescent monoclonal cells were isolated to 48-well plates and sub-cultured to 12-wells plates. Confluent monoclonal GFP+ cells colonies were harvested, in which 4/5 cells were cryopreserved and other were lysed for PCR amplification to identify the integration of transgene **(Supplementary Table S3)**.

### 2.5. Generation of cloned pigs by somatic cell nuclear transfer

The positive transgenic cells were treated with Cre enzyme (Excellgen, Rockville, MD USA) to delete the EGFP marker gene before somatic cell nuclear transfer (SCNT). Approximately 200 reconstructed embryos were surgically transferred to the oviduct of recipient gilts 24 hours after estrus was observed. The pregnancy status was detected using an ultrasound scanner at 26 day after surgery. This pregnancy status was then monitored monthly from the embryo transfer to birth. All the cloned piglets were naturally born.

### 2.6. PCR and southern blotting analysis of founder pigs

Genomic DNA from ear tissue of founders (F0) pigs was isolated according to the protocol of the OMEGA Kit (OMEGA Bio-Tek, Georgia, USA) for PCR and southern blot analysis. PCR was performed using primers shown in **Supplementary Table S3**. The products were resolved by 1 % agarose gel electrophoresis, and sequencing to determine the occurrence of KI.

Southern blot analysis were performed with 20 μg of genome DNA, after digested with restriction enzyme, The digested DNA were extracted with ethanol precipitation, and separated in a 0.8 % agarose gel, and then transferred to a nylon membrane. The membrane was hybridized with digoxigenin-labeled DNA probe which targets BgEgXyAp gene. Hybridization and washing were performed according to the procedures of DIG-High Prime DNA Labeling and Detection Starter Kit II (Roche, Mannhein, Germany). The membranes were imaged using UVP software.

### 2.7. Gene copies quantification

The PB-PSP-BEXA plasmid was diluted to different concentration (copies: 5^0^, 5^1^, 5^2^, 5^3^, 5^4^, 5^5^) as a standard curve, and simultaneously subjected to real time PCR amplification with 10 ng genomic DNA of the KI pigs (see the prime RTQ-copy in **Supplementary Table S3**). Standard curve was generated according to the standard product copies and CT value, and the corresponding copy number of each sample was calculated per standard curve.

### 2.8. Enzymatic activity assay and western blot

The saliva was collected from growing KI pigs and age-matched non-KI pigs. To measure the saliva enzymatic activity of transgene, 3, 5-dinitrosalicylic acid (DNS) method was empolyed to determine β-glucanase and xylanase activity, as described previously (Luo *et al*. 2010; Liu *et al*. 2010; Zhang *et al*. 2018). One unit of activity was defined as the quantity of enzyme that releases reducing sugar at the rate of 1 μmol/min. To detecte phytase activity, method of vanadium molybdenum yellow spectrophotometry was used as described previously (Zhang *et al*. 2018). One unit of phytase activity was defined as the amount of activity that liberates one micromole of phosphate per minute at 39°C.

For western blot, total proteins from saliva was ultrafiltrated using an amicon ultra 15 mL centrifugal filter (Millipore, Massachusetts, USA), 20 μg of saliva total protein were subjected to SDS polyacrylamide gel electrophoresis, followed by immunoblotting onto a polyvinylidene fluoride membrane (Millipore, Massachusetts, USA). Membranes were blocked with 5% non-fat dry milk in TBST for 2 hours and incubated overnight at 4°C with primary rabbit polyclonal antibodies against BG17A and EAPPA (Zhang *et al*. 2018). The blots were probed with a salivary amylase antibody (ab34797, Abcam) to confirm equal protein loading. After the membranes washed with TBST, they were further incubated with a secondary HRP-conjugated goat anti-rabbit IgG antibody for 2 hours at room temperature and imaged using a supersignal west pico enhanced chemiluminescence kit (Thermo Fisher Scientific, Suwanee, GA,USA). The signals were visualized with UVP software.

### 2.9. Statistical analysis

All results data were presented as mean±SEM. To assess the statistical significance of differences, data were analyzed using One-way ANOVA followed by Duncan’s test. P < 0.05 was considered statistically significant.

## 3. Result

### 3.1. Screening of sgRNA with high cleavage efficiency in CEP112

We designed 7 sgRNAs targeting intron 5 region of the porcine *CEP112* gene. We first co-transfected CRISPR-targeting plasmid and fluorescence expressing plasmid (pBb-ubc-eGFP) to evaluate the transfection efficiency **(Fig.1 a)**. The cleavage efficiency of each targeting plasmid was evaluated with T7EI assays. The results demonstrated that some sgRNAs can efficiently induced DSBs in targeted loci, such as C1, C3, C5 and C7 had more than 40% cleavage efficicency, but some sgRNAs also showed lower efficiency **(Fig.1 b)**. To further confirm the cleavage rates of the sgRNAs, C3, C5 and C7 PCR products were cloned into TA vector, and sequenced the assess the percentage of mutant alleles. The results showed that the rates of presence of indels in cleavage sites were 27.8%, 36.7% and 23.0% for target loci of C3, C5 and C7, respectively **(Fig.1 c)**. We selected C5 sgRNA with the highest cleavage rate in the subsequent experiments.

**Figure 1.**
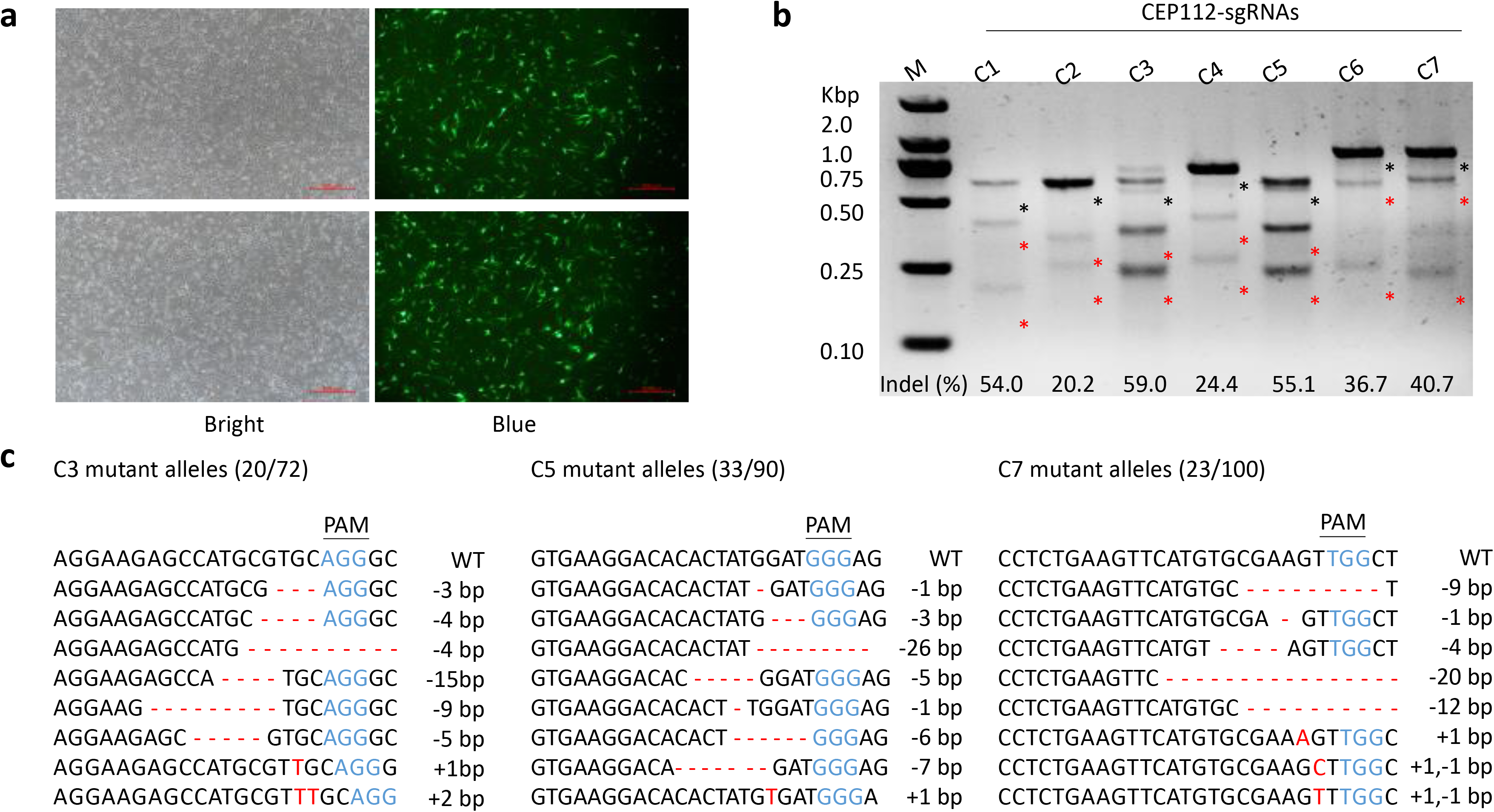
Screening of sgRNA with high cleavage efficiency in *CEP112* loci. (a) Fluorescence expression in PFFs co-transfected with sgRNAs and pBb-ubc-eGFP. (b) Determination of the cleavage efficiency by T7E1 assay. The red and black asterisks indicate cut and uncut bands, respectively. (c) Sanger sequencing further confirmed the cleavage rates of sgRNAs of C3, C5 and C7 in the *CEP112* locus. Eight mutant alleles were presented as representative. Indels were shown in red letters and dashes. PAM sequences were shown in blue.

### 3.2. Optimization of the ratio of homologous lengths

To investigate the effect of homologous length on the KI efficiency of large fragment (≈ 20 kbp) into porcine genome region, we constructed 6 donor templates of *ROSA26* locus and 4 templates of *CEP12* locusm, which contain combinations of homologous arms (HA) with different sizes. The primer sets were used to amplify HA for identifying the KI cell lines, locating at genome and donor vector regions adjacent to the homology arm ends **(Fig.2 a)**. Fluorescent monoclonal cells **(Fig.2 b)** were picked up and genotyped by PCR assay and agarose gel electrophoresis **(Fig.2 c)**. For the *ROSA26* locus, the donor plasmid ROSA26-LA320RA3769 had the highest HDR efficiency (7.94%), but equal length HA could not integrate large fragment into the pig genome in CEP112???. Furthermore, when LA was shorter (< 500 bp) or the ratio between LA and RA increased, the KI efficiency became higher **(Table.1)**. The donor vector (CEP112-LA340RA3219) contained LA of 250 bp and RA of 3219 bp generated up to 13.89% CRISPR/Cas9-mediated HDR efficiency, significantly higher than *ROSA26* **(Table.1)**. Furthermore, we investigated whether different structure of *CEP112* donor can influence the KI efficiency. We found that the KI efficiency was not different between supercoiled plasmidand linearized plasmid, **(Table.1)**, which indicated that using simple supercoiled plasmid can achieve the equivalent KI rate as linearized plasmid.

**Figure 2.**
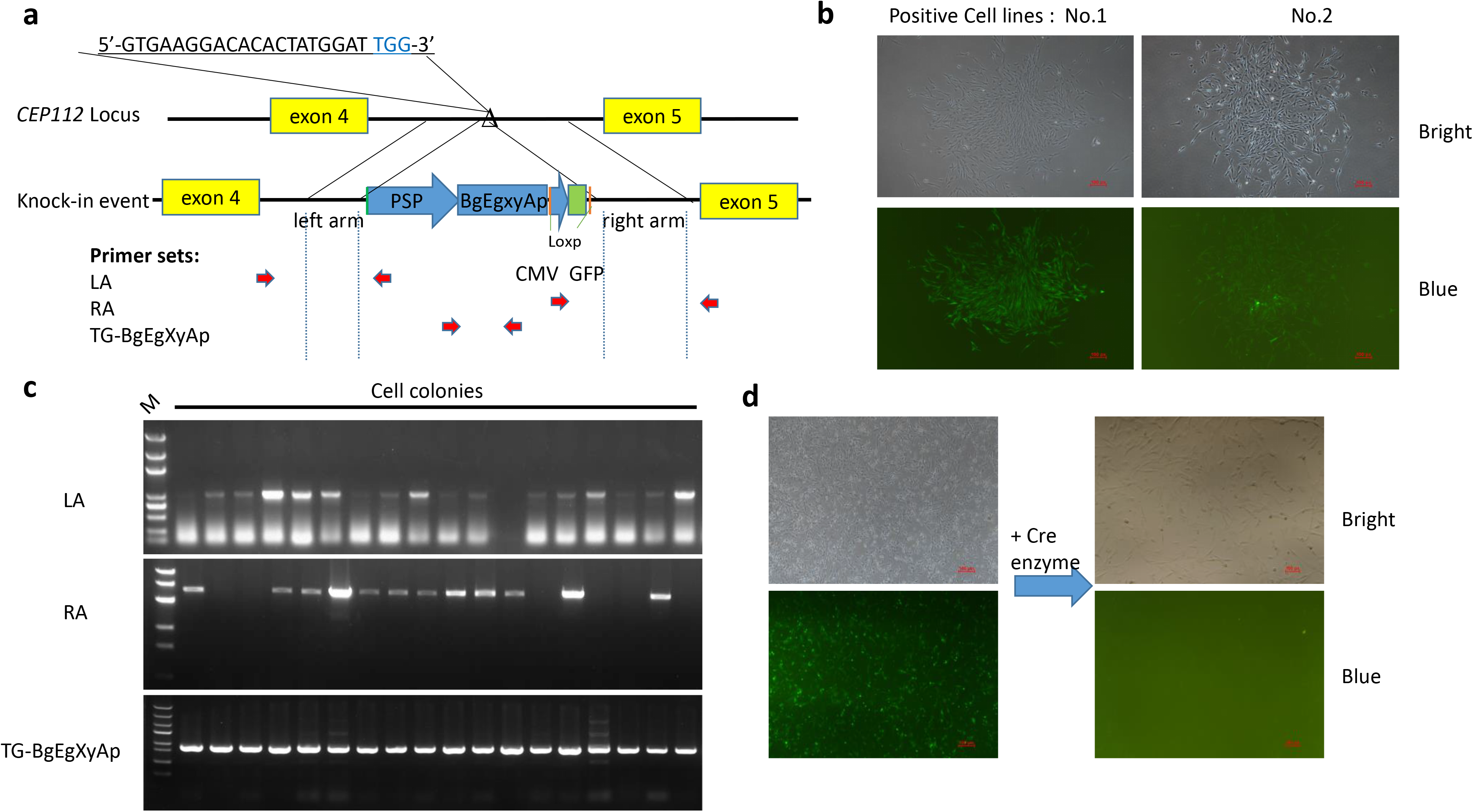
Effects of the length of HA on the screening of KI cell lines. (a) Schematic diagram of the strategy for insertion BgEgXyAp gene and fluorescence labeling gene into porcine *CEP112* intron 5. Triangles indicate Cas9 targeting sites, blue letters is PAM sequences, Pink dashes indicate loxp sequence and red arrows are primer sets for PCR assay. (b) Fluorescent monoclonal cells screening. (c) Analysis of knock-in event by gel electrophoresis. (d) Fluorescent labeling gene was deleted by cyclization recombination (Cre) enzyme before SCNT.

**Table 1.** Summary of KI events of different proportions of homology arms

|  | plasmid<br>(LARA)<sup>a</sup> | Target events |  |  |  |  |
| --- | --- | --- | --- | --- | --- | --- |
|  |  | LA | RA | BgEgxyAp | HDR efficiency<sup>b</sup> | LA/RA |
| **Supercoiled** | CEP112-LA1199RA1999 | 0/30 | 0/30 | 0/30 | 0% | 1:2 |
|  | CEP112-LA2003RA1999 | 0/50 | 0/50 | 0/50 | 0% | 1:1 |
|  | CEP112-LA250RA3219 | 11/72 | 10/72 | 11/72 | 13.89% | 1:13 |
|  | CEP112-LA340RA3219 | 30/201 | 26/201 | 26/201 | 12.94% | 1:9 |
|  | ROSA26-LA320RA1214 | 0/29 | 0/29 | 0/29 | 0% | 1:4 |
|  | ROSA26-LA320RA2046 | 0/31 | 0/31 | 0/31 | 0% | 1:6 |
|  | ROSA26-LA320RA3769 | 5/63 | 6/63 | 5/63 | 7.94% | 1:12 |
|  | ROSA26-LA1040RA1214 | 0/27 | 0/27 | 0/27 | 0% | 1:1 |
|  | ROSA26-LA1040RA2046 | 3/38 | 2/38 | 3/38 | 5.26% | 1:2 |
|  | ROSA26-LA1040RA3769 | 4/72 | 2/72 | 2/72 | 2.78% | 1:3 |
| **Linearized** | CEP112-LA250RA3219 | 7/132 | 9/132 | 7/132 | 5.30% | 1:13 |
|  | CEP112-LA340RA3219 | 11/72 | 9/72 | 9/72 | 12.50% | 1:9 |

<sup>a</sup> Donor plasmid, for example CEP112-LA1199RA1999 represents with 1199bp LA and 1999bp RA .
<sup>b</sup> HDR efficiency (%) = The minimum events in LA, RA and BgEgxyAp / total cell lines \* 100% .

### 3.3. Generation of BgEgXyAp KI pigs

Total three cell lines (*CEP112* locus KI) which exhibited a better morphology and viability were carefully singled out. After fluorescent labeling gene deletion **(Fig.2 d)**, cells were pooled and used as nuclear donors for somatic cell nuclear transfer. We transferred total of 876 reconstructed embryos into 4 recipient gilts. Two recipients were pregnant and delivered 9 Duroc piglets (Table 2). PCR screening results showed that 4 heterozygous (503, 605, 705 and 711) achieved site-specific KI in one allele of *CEP112* locus **(Fig.3 a)**, but only one male piglet (705) was alive **(Fig.3 b)**. Sequencing also showed that donor sequence could be detected on both the 3’-end of LA and the 5’-end of RA without excessive mutations in all KI pigs **(Fig.3 c)**. However, the other *CEP112* allele were KO with varied insertions or deletions in KI pigs, including thymine(T) to cytosine (C) mutation (605 and 705) or 11 bp deletion (503 and 711). Non-KI pigs showed homozygous modification with 11 bp deletion (501), 100 bp deletion (603) or T to C mutation (601 and 709), and biallelic modification with 18 bp insertion on one allele and 20 bp deletion on the other allele (703), or 2bp deletion on one allele and adenine to cytosine mutation on the other allele (701) **(Fig.3 d)**. Among the piglets, 2 were stillborn (1 KI piglet and 1 non-KI piglet), 5 were born weak and died within 1 month (2 KI piglet and 3 non-KI piglets), and 3 appeared normal at birth and survived to date (1 KI pig and 2 non-KI pigs) **(Table 2)**.

**Table 2.**
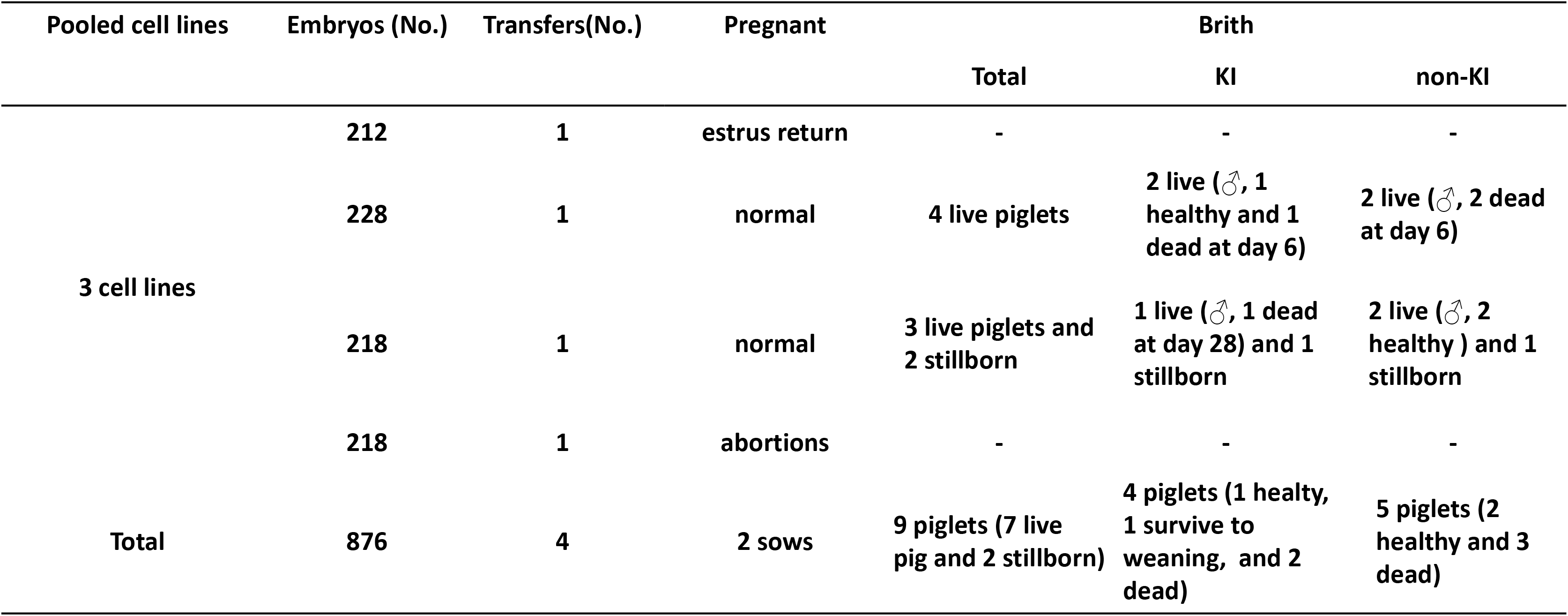
Embryo transfer data for cloned pigs

| Pooled cell lines | Embryos (No.) | Transfers(No.) | Pregnant | Brith |  |  |
| --- | --- | --- | --- | --- | --- | --- |
|  |  |  |  | Total | KI | non-KI |
| 3 cell lines | 212 | 1 | estrus return | - | - | - |
|  | 228 | 1 | normal | 4 live piglets | 2 live (♂, 1 healthy and 1 dead at day 6) | 2 live (♂, 2 dead at day 6) |
|  | 218 | 1 | normal | 3 live piglets and 2 stillborn | 1 live (♂, 1 dead at day 28) and 1 stillborn | 2 live (♂, 2 healthy ) and 1 stillborn |
|  | 218 | 1 | abortions | - | - | - |
| Total | 876 | 4 | 2 sows | 9 piglets (7 live pig and 2 stillborn) | 4 piglets (1 healty, 1 survive to weaning, and 2 dead) | 5 piglets (2 healthy and 3 dead) |

**Figure 3.**
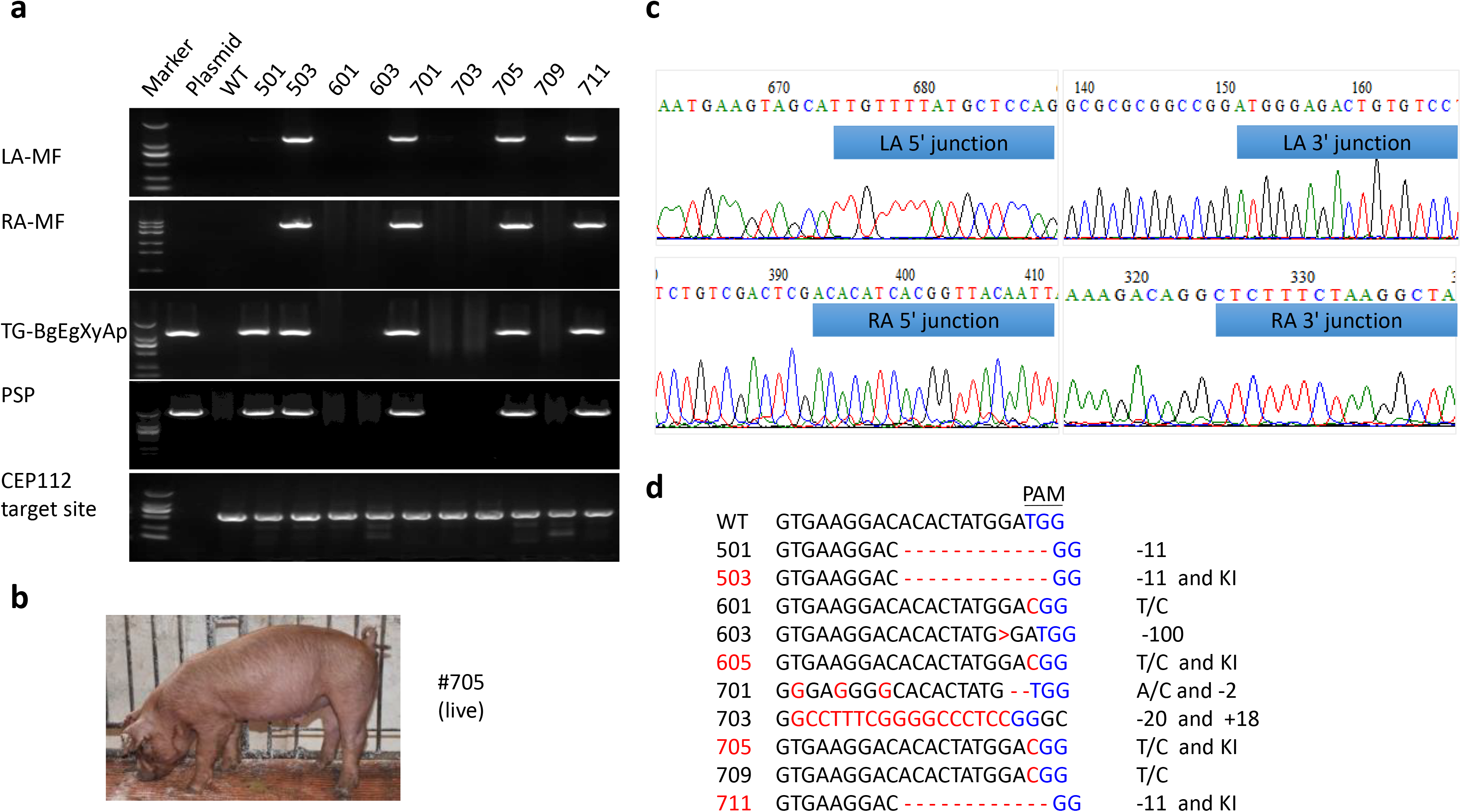
Generation of cloned pigs using CRISPR/Cas9 mediated HDR system. (a) Genomic DNA of the cloned pigs were amplified by PCR and analyzed by gel electrophoresis. (b) The live KI pigs (705) aged 1 months. (c) PCR products of KI loci were sequenced. Blue represents the junction sequences of the homology arms and KI locus. (d) Sequences of targeted *CEP112* loci of all cloned pigs. KI pigs are showed with red number. Red letters and dashes represented as indels. PAM sequence was shown in blue.

### 3.4. Characterization of gene integration copies and expression patterns in modified pigs

The plasmid was diluted to different copies to generate standard curve to calculate the copy numbers of transgene with real time PCR. The results showed that all modified pigs have approximately one copy, implying a precise KI without random integration **(Fig.4 a)**. Furthermore, southern blot also confirmed that only single copy was inserted into *CEP112* locus by using two kinds of restriction endonucleases *BsrGI* and XmnI, respectively **(Fig.4 b)**. During feeding period, we collected the saliva from KI and non-KI pigs at 1 month and 6 month for enzymatic activity assays. The results showed that the enzyme activity of KI pig (705) was increased with age, and when 6 months old, saliva can produce 5.49 U/mL of β-glucanase, 1.42 U/mL of xylanase, and 0.77 U/mL phytase **(Fig.4 c)**. Western blot also confirmed the presence of protein of β-glucanase (BG17A), EAPPA (phytase) in the saliva of modified founders **(Fig.4 d)**.

**Figure 4.**
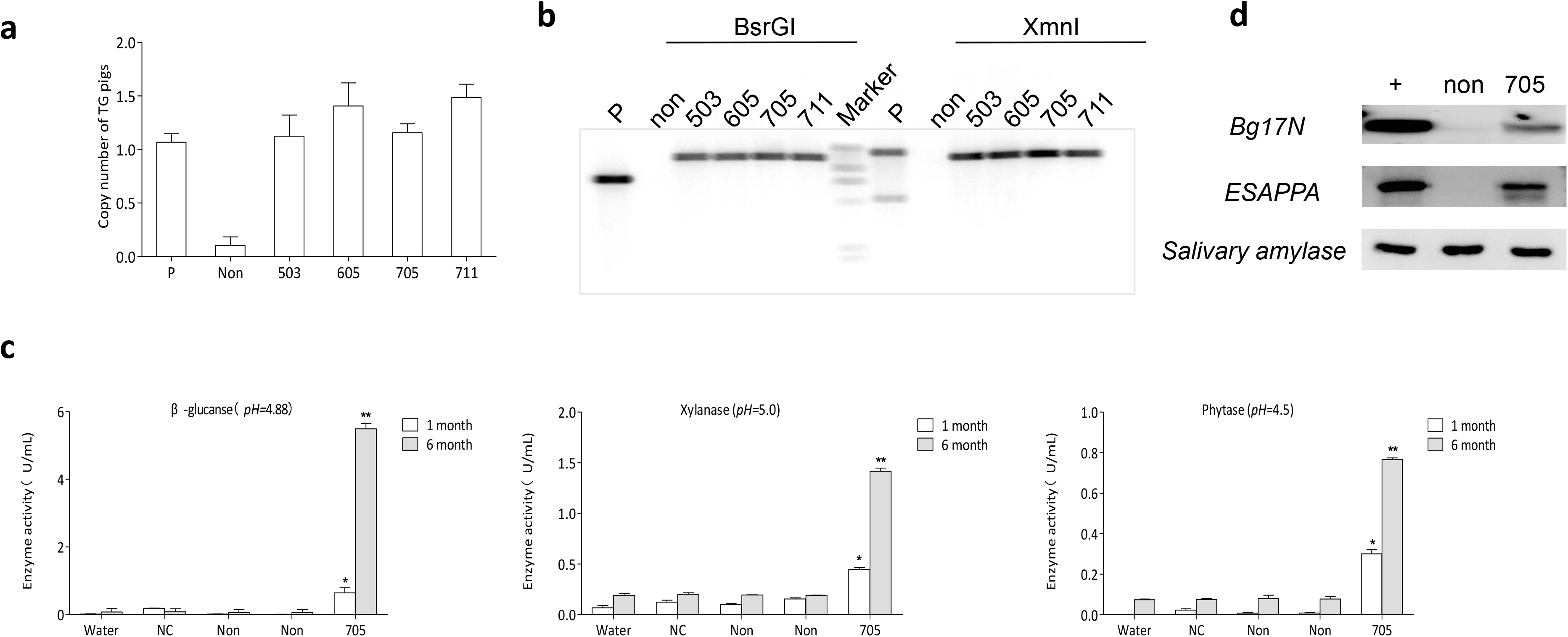
Presence of transgene and expression in the KI pigs. (a) Calculation of copy number of integrated gene in KI pigs according to the standard curve. (b) Southern blot analysis result of multi-enzyme transgene integration in KI pigs. Genomic DNA of KI pig was digested by two restriction endonucleases *BsrG*I and *Xmn*I, respectively. (c) Salivary β-glucanase, xylanase and phytase activity assays of KI pigs at 1 month and 6 month. (d) Western blotting demonstrated the expression of Bg17N and EAPPA in the saliva of KI pig (705). Plus sign indicates positive sample; P represents donor vector; NC and non represent non-modified pig and non-KI pig, respectively. Values are shown as mean ± SEM. The asterisk represents P < 0.01.

## 4. Discussion

Previous report on gene targeting study showed a correlation between the length of HA and KI efficiency (0.2-14 kbp) (Thomas and Capecchi 1987). When HA was less than 200 bp, the HDR efficiency would be significantly decreased, and the length reached about 14 kbp or more, the relationship gradually became insignificant. However, in the presence of artificial nuclease induced DSBs, KI efficiency can be improved greatly with successful gene editing from HA lengths ranging from 9 bp to several 5 kbp (Orlando *et al*. 2010; Shin *et al*. 2014; Byrne *et al*. 2015). Although numerous studies have exploited engineered nucleases to develop multiple targeting strategies and various genomic modifications have been achieved, the integration fragments were shorter than <10 kbp (Chu *et al*. 2015; Sakuma *et al*. 2015; Sakuma *et al*. 2016; Song *et al*. 2016; Maruyama *et al*. 2016; Nakamae *et al*. 2017). Thus, precise and efficient genetic modification with large fragment KI (> 10 kbp) remains difficult in primary fibroblasts. In this study, we found a combination of HAs about 300 bp (LA) and 3 kbp (RA) can yield an efficient KI with an exceeding 10% efficiency (*CEP112* loci). Interestingly, previous reports also found that shorter LA and longer RA can efficiently enhance CRISPR-mediated integration (Shin *et al*. 2014; Zhang *et al*. 2017). The optimal HA system was very similar to the DNA length dependence DSB repair such as single-strand annealing (SSA) pathway in RA or MMEJ pathway in LA (more frequent than HDR pathway) (Ceccaldi *et al*. 2016), suggesting that our system was probably repaired by the MMEJ or SSA pathway. Furthermore, our results shown the KI efficiency was insignificant when the gRNA target sequence was introduced into the LA 5’ end compared to the linearized plasmid, which seems inconsistent with previous studies (Bollag *et al*. 1989). Bollag et al found that linearized donor vectors after enzyme restriction significantly increased HDR efficiency (Bollag *et al*. 1989), and Shin et al also confirmed that linear donor was easier to achieve site-specific integration (Shin *et al*. 2014). We speculated that gRNA can efficiently induce DSBs in donor plasmid after transfection and generate the same structure as linearized plasmid. Meanwhile, the structure of donor template may not be essential for the CRIPSR-mediated large fragments integration. In generally, we reasoned the use of shorter LA and longer RA can make the precise modification more feasible and accessible.

In order to prevent the abnormal karyotype induced by gene modification from affecting the normal development of reconstructed embryos, resulting in low birth rate of cloned pigs, we mixed multiple positive clones as nuclear donors for somatic cell nuclear transfer. A total of 876 reconstituted embryos were constructed and transplanted into surrogate recipients, 2 of which were pregnant and gave birth finally. A total of 9 cloned piglets were produced, in which 5 piglets were identified as negative for transgene. The main reason may be that the positive clones are impure and mixed with some wild-type cells, which cannot be distinguished by PCR for KI manipulation. In addition, among the 9 cloned pigs, two were stillborn and seven were alive. Among the 7 live piglets, 1 KI piglets and 2 non-KI piglets died within 1 week after birth, 1 KI pig lived to the weaning stage, and only 1 KI pig and 2 non-KI cloned pigs remained healthy. Previous studies have reported abnormal reprogramming of cloned embryos or incomplete embryonic development during pregnancy could account for the early death of cloned animals (Carter *et al*. 2002). Therefore, cloned pigs have high deformity rate and mortality produced by SCNT, and the loss of cloned piglets is generally high (58%) before weaning (Liu *et al*. 2015; Schmidt *et al*. 2015). Our results also showed a similar mortality rate (KI 50% (2/4), non-KI 40% (2/5)) as previous reports (Liu *et al*. 2015; Schmidt *et al*. 2015).

Traditionally, targeted insertions into the *ROSA26* or *H11* locus are frequently used for the constitutive or conditional expression of transgenic pigs (Jakobsen *et al*. 2011; Li *et al*. 2014; Lin *et al*. 2014). Here we established a CRISPR/Cas9 mediated approach for generating KI pigs in *CEP112* locus, a potential safe harbor in the pig genome. The results shown that *CEP112* locus supports exogenous genes expression driven by a tissue-specific promoter like other known safe harbors, *ROSA26* (Li *et al*. 2014; Ruan *et al*. 2015; Xie *et al*. 2017). Meanwhile, we observed that targeting *CEP112* were more efficient for gene insertion than *ROSA26* using our optimal HA system. We guessed that it was mainly caused by the chromatin structure, suggesting that the *CEP112* locus may be in an open genomic region and serves as a “hot spot” for gene editing. Many studies have found that KI event was more likely to occur in the regions where DNA can be frequently duplicated or transcribed (Frohman and Martin. 1989; Ruan *et al*. 2014).

In summary, we achieved the site-specific targeting of large fragment in *CEP112* locus in pig somatic cells using our optimal HA system, which have the potential to serve as a platform for effective generation of precisely modified pigs for biomedical and agricultural applications. Finally, we also expected that the optimal system can be adopted and further strengthen the notion that *CEP112* is a safe locus for gene addition.

## Acknowledgements

This work was supported by grants from the National Science and Technology Major Project for Breeding of New Transgenic Organisms (2016ZX08006002).

